# Latent tuberculosis infection in foreign-born communities: import *vs*. transmission in the Netherlands derived through mathematical modelling

**DOI:** 10.1101/228924

**Authors:** Hester Korthals Altes, Serieke Kloet, Frank Cobelens, Martin Bootsma

## Abstract

While tuberculosis represents a significant disease burden worldwide, low-incidence countries strive to reach the WHO target of elimination by 2025. Screening for TB in immigrants is an important component of the strategy to reduce the TB burden in low-incidence settings. An important option is the screening and preventive treatment of latent tuberculosis infection (LTBI). Whether this policy is worthwhile depends on the extent of transmission within the country, and introduction of new cases through import. Mathematical transmission models of tuberculosis have been used to identify key parameters in the epidemiology of TB and estimate transmission rates. An important application has also been to investigate the consequences of policy scenarios.

Here, we formulate a mathematical model for TB transmission within the Netherlands to estimate the size of the pool of latent infections, and to determine the share of importation –either through immigration or travel- versus transmission within the Netherlands. We take into account importation of infections due to immigration, and travel to the country of origin, focusing on the three ethnicities most represented among TB cases, excluding those overrepresented in asylum seekers: Moroccans, Turkish and Indonesians. We fit a system of ordinary differential equations to the data from the Netherlands Tuberculosis Registry on (extra-)pulmonary TB cases from 1995-2013.

We find that for all three foreign-born communities, immigration is the most important source of LTBI, but the extent of within-country transmission is much lower (about half) for the Turkish and Indonesian communities than for the Moroccan. This would imply that contact investigation would have a greater yield in the latter community than in the former. Travel remains a minor factor contributing LTBI, suggesting that targeting returning travelers might be less effective at preventing LTBI than immigrants upon entry in the country.

## Introduction

Revised estimates of the burden of latent tuberculosis infection (LTBI) indicate that about a quarter of the world population is infected with tuberculosis (TB) (1). Recently, guidelines for the management of LTBI were formulated against the background of the World Health Organization’s (WHO) End TB Strategy (2, 3). With the post-2025 targets of TB elimination, even low-incidence countries are seeking to decrease TB incidence. Some of these countries have already implemented screening for latent TB infection at entry in the country, others are envisaging implementing this measure. The Netherlands, for example, is starting pilots among different immigrant groups to evaluate feasibility of LTBI screening at entry. For the critical assessment of LTBI screening policies, it is necessary to quantify the extent to which screening at entry is better than searching for infections in the context of contact investigation. Essentially, we wish to know what fraction of new LTBI in immigrants results from importation of infections, from transmission within the country, or from travel to the country of origin while living in the Netherlands. Cohort studies have provided estimates for the percentage of immigrants with a positive interferon-gamma release assay (IGRA) test at entry (4). However, the coverage of contact investigation in foreign-born groups is poor (5) and estimates of LTBI in this context are therefore incomplete.

Mathematical transmission models of tuberculosis have been used to identify key parameters in the epidemiology of TB (6) and estimate transmission rates (7). An important application has also been to investigate the consequences of policy scenarios (1, 8, 9). Here, we formulate a mathematical model for TB transmission within the Netherlands to estimate the size of the pool of latent infections, and to determine the share of importation -through either immigration or travel- versus transmission within the Netherlands. We take into account importation of infections due to immigration, and travel to the country of origin, focusing on the three ethnicities most represented among TB cases, excluding those overrepresented in asylum seekers. To that end, we fitted a system of ordinary differential equations to the data from the Netherlands Tuberculosis Registry on (extra-)pulmonary TB cases from 1995-2013, to estimate the TB transmission parameter, as well as the relative fraction of recent versus remote LTBI at the start of the study period for the three ethnic groups considered.

## Methods

We formulate a system of ordinary differential equations that is as realistic as possible, yet simple enough to allow the fit to time series data, i.e., the number of unknown parameters is limited. First, we focused our model on the 1^st^ generation immigrants, to avoid the need to describe the birth process in the immigrant population leading to 2^nd^ generation immigrants, and the complexities associated with mixed marriages. For all three ethnicities considered, the number of 2^nd^ generation cases from non-mixed marriages represent about 5% of cases from the country in question (Appendix, Table A1).

Second, we assumed that TB transmission in the Netherlands occurs within the ethnic groups. There is no recent literature on this matter, but twenty years ago, Borgdorff *et al.* (10) estimated from TB typing data spanning a 2-year period that the extent of cross-ethnicity transmission was ethnicity-dependent. Most Turkish cases result from transmission within the ethnic group, while a third of recently transmitted TB in Moroccans was due to infection from individuals from other nationalities (10).

We simplify it for the sake of the fitting procedure and will consider implications of this simplification in the discussion.

We first present the deterministic model describing the transmission and pathogenesis of TB in the Netherlands, calibrating it on data from the National Tuberculosis Registry (NTR) and Statistics Netherlands (11). We exclude countries of origin from which most immigrants are asylum seekers, as they are associated with a complex immigration and screening process, which requires a different mathematical model. We consider Moroccan (M), Turkish (T) and Indonesian (I) first generation immigrants (FGIs) in the Netherlands. These ethnic groups had the highest cumulative contribution to TB cases in the Netherlands over the last two decades (12, 13). Because of changing immigration and emigration over the study period (14, 15), we do not assume a fixed population size.

### Data sources

The National Tuberculosis Registry (NTR) provided the data on all TB cases notified in The Netherlands between 1993 and 2013 (16). Variables included were: date of diagnosis, disease site (pulmonary/extra-pulmonary/both), country of birth, ethnicity, time spent in the Netherlands at time of diagnosis, reason for testing (contact-tracing/screening/symptoms etc), immigrant status (immigrant/asylumseeker etc), and time from symptoms to diagnosis.

To minimize the effect of underreporting and incomplete patient files due to running cases diagnosed before 1993, our research studies the years 1995 up to and including 2013. Also, (monthly) immigration data by ethnicity, from Statistics Netherlands (14), was only available starting in 1995. Prevalence of TB in the country of origin was obtained from the WHO TB database (17, 18)(Appendix, Table A2).

### SEIS model

Since compartment sizes and a number of parameter values differ between ethnic groups, we use subscript *k* as a label for the country of origin. Table 1 indicates whether model parameters are country-specific, or the same across ethnic groups.

**Table 1.**
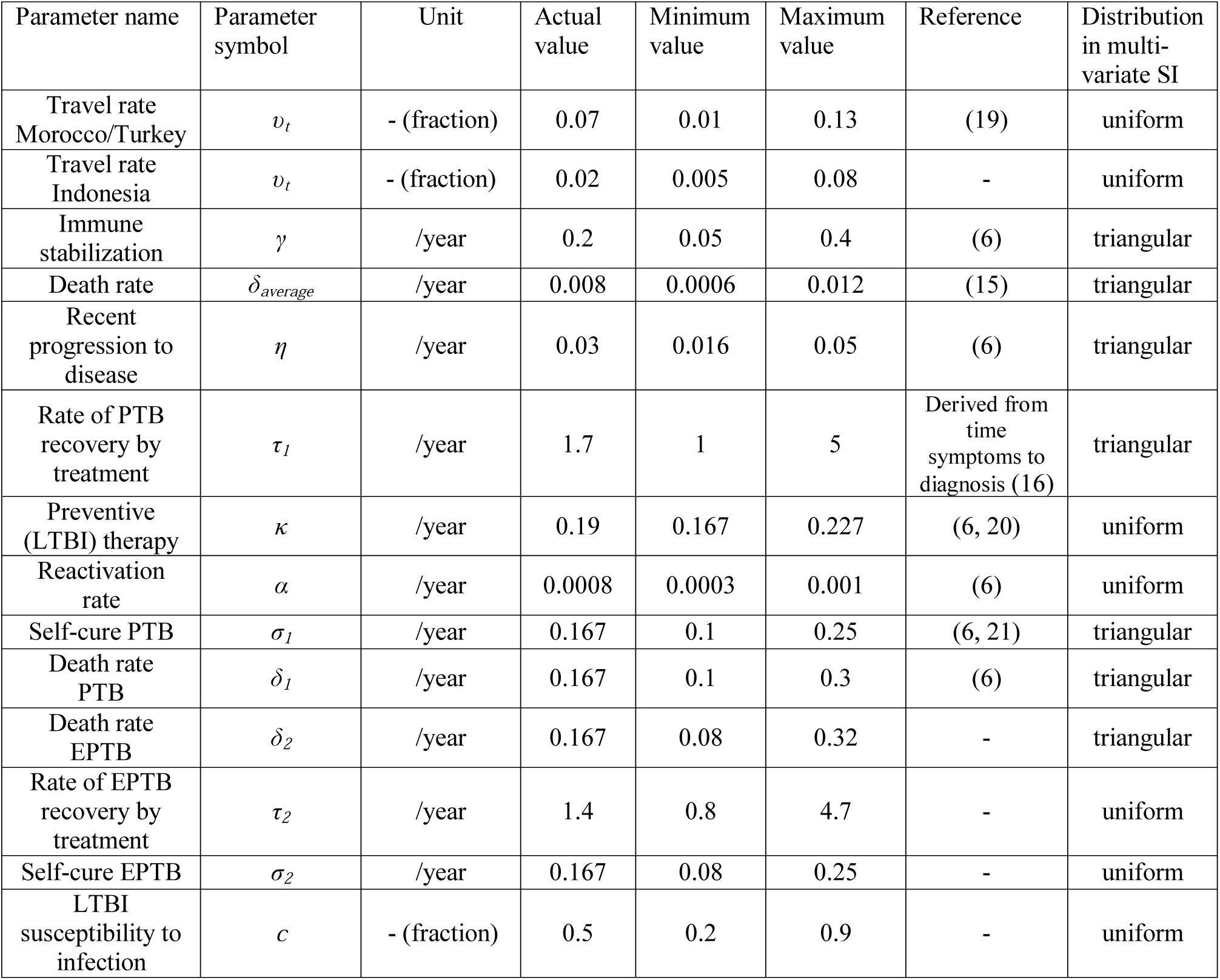
Model parameters

### Model description

To determine the relative impact of immigration and travel on new LTBIs we developed a compartmental SEIS model for the three ethnic groups (Fig 1; mathematical formulation in appendix, Section A1). It is adapted from previous SEIR-models of TB transmission (6).

**Fig 1.**
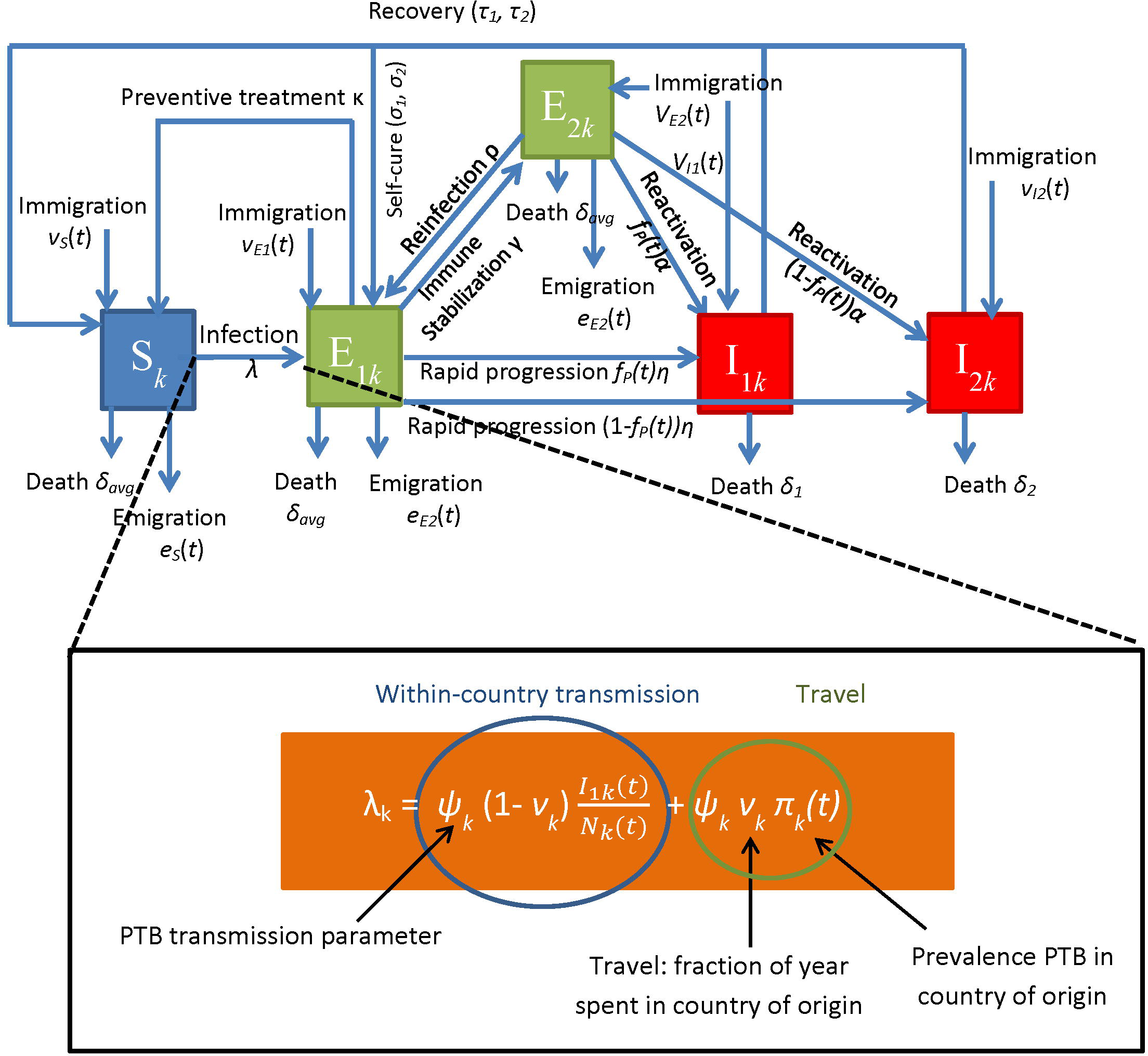
Schematic of ODE model.

The population is divided into 5 compartments: individuals susceptible to TB infection (*S*), recent and remote LTBI (*E*_1_ and *E*_2_ respectively), and pulmonary TB cases (PTB) and extra-pulmonary (EPTB) cases (*I*_1_ and *I*_2_ respectively). When we refer to PTB cases, we implicitly include cases with both extra-pulmonary and pulmonary TB, as they are also infectious–while EPTB cases are not.

The inflow into the susceptible compartment (*S*) of foreign-born from country *k* consists of immigrants (*v_s_(t)*), cases cured from TB infection through preventive treatment of recent LTBI (*κE_1_*), recovery through treatment (*τ_1_I_1_* and *τ_2_I_2_*) of infectious TB cases. Susceptibles may die at rate *δS*, emigrate at rate *w_S_(t)* or get infected at rate *λ_k_S*, with *λ_k_* the force of infection which we will specify later. The emigration rate *w_S_(t)* is the yearly emigration number (appendix Table A3) weighed with the share of susceptibles among non-diseased individuals, see appendix section A1 for the formula. Upon infection, a susceptible establishes a recent latent tuberculosis infection (LTBI). The recent LTBI compartment (*E*_1_) can also grow due to reinfection of remote LTBI, at rate c*λ_k_E_2_*, or due to immigrants entering the Netherlands with recent LTBI (*v_El_*). Also, PTB and EPTB cases may self-cure (at rate *σ_1_I_1_* and *σ_2_I_2_* respectively) and become recent LTBI. Individuals leave the *E_1_*-compartment because of disease progression at rate *ηE_1_*, they may turn into a remote LTBI (at rate *γE*_1_) or die at rate *δE_1_* or emigrate at rate *W_*E*1_(t)*. Emigration of recent LTBI is again the yearly emigration rate (appendix, Table A3) weighed with the share of recent LTBI. The inflow into the remote LTBI compartment (*E_2_*) further consists of immigration at rate *v_E2_*; remote LTBI’s disappear by emigration at rate *w_E2_(t)* (emigration weighed with share of remote LTBI) and through death (*δE_2_*) as in the *S*- and *E_1_*-compartment. They may get reinfected at rate *cλE_2_* to become recent LTBI. Through reactivation, a remote LTBI turns into active TB at rate *αE_2_*. The *I*-compartments refer to TB cases, pulmonary and infectious (PTB, *I_1_*) or extra-pulmonary (EPTB, *I_2_*). The inflow in these compartments consists of rapidly progressing LTBI cases at rate *f_P_(t)ηE_1_* and [1-*f_P_(t)*]*ηE_1_* in *I_1_* and *I_2_*Irrespectively, where *fp(t)* is the fraction PTB (or P+E) among all TB cases over the study period. This fraction may change as a linear function of time a *f_p_(t)= f_p0_-bt* over the study period, where *f_p0_* is the fraction PTB over all TB cases in 1994-1995. Thus, the fraction LTBI developing EPTB vs PTB may change, due to a changing population composition for example. Reactivation of remote LTBI yields *I_1_* cases at rate *f_p_(t)αE_2_*, and *I_2_* cases at rate [1-*f_p_(t)*]*αE_2_*. We assume that infectious TB cases do not emigrate. PTB cases die at rate *δ_1_I_1_* and EPTB cases at rate *δ_2_I_2_*. PTB and EPTB cases recover through self-cure at rate *σ_1_I_1_* and *σ_2_I_2_* respectively, and through treatment at rate *τ_1_I_1_* and *τ_2_I_2_* respectively.

To describe the force of infection, we need *Ψ_k_*, the TB transmission parameter, which depends on mixing rate and other contact patterns. We assumed it to be the same within an ethnic group in the Netherlands and in the country of origin. In the sensitivity analysis we will test the impact on results of closer contact rates in the country of origin compared to the Netherlands. The parameter *v_k_* is the fraction of time a susceptible immigrant spends in country of origin. *N* is the total immigrant population size and *π_k_(t)* is the prevalence of PTB in country of origin.

The force of infection is mathematically captured as follows:

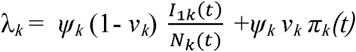

The first term of the force of infection, 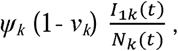 represents transmission of TB in the Netherlands.

The second term, *Ψ_k_ v_k_ π_k_(t)*, represents TB acquisition when traveling to the country of origin. We refer to these terms as the ‘transmission term’ and the ‘travel term’ respectively.

### Travel rate and immigrant TB

The ‘travel term’ consists of the TB transmission parameter *Ψ_k_*, the travel rate *v_k_* and the prevalence in country of origin *π_k_(t)*. The travel rates for the Moroccan and Turkish communities were derived from a study by Kik *et al.* in Rotterdam communities (19). The derivation of the travel rate term is shown in the appendix (section A2).

Import of LTBI, remote and recent, was estimated as follows:

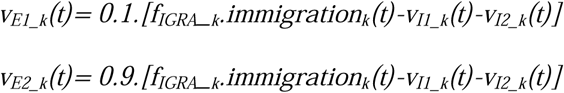

where *f_IGRA_k_* is the proportion of immigrants entering the Netherlands from country *k*, having a positive IGRA (Interferon Gamma Release Assay) test - as a proxy for LTBI. This assumed that 10% of non-infectious IGRA+ individuals have recent LTBI while 90% have remote LTBI. Data is shown in appendix (Table A3).

The number of new (E)PTB cases directly imported were considered those in the NTR recorded as having entered the Netherlands less than 6 months prior to diagnosis (Appendix, Table A4).

### Maximum likelihood

We applied a maximum likelihood (ML) method to find the best fit for our target parameters *b,Ψ* and *r_E1_* similar to the method described in Diekmann *et al*. (22). Details of the fitting procedure are given in the appendix (Section A3).

### Sensitivity analysis

#### Univariate sensitivity analysis

This was carried out by fitting the ODE model to the NTR and CBS data, varying one parameter at a time, setting to its minimum or maximum value (Table 1), while keeping the other parameters at their default value (Table 1, “Actual”).

#### Multivariate sensitivity analysis

95% intervals on the estimates for TB transmission, fraction initial remote versus recent LTBI and percentage contribution to LTBI were derived with a multivariate analysis. This involved drawing the 14 parameters in Table 1, as well as the parameter governing the fraction of LTBI becoming EPTB compared to PTB (*b*), from a uniform or triangular distribution with minima and maxima as indicated in Table 1. With each combination of these parameters drawn, the best-fit model was derived with its estimate for the transmission rate parameter, rate of change in fraction LTBI becoming PTB vs EPTB, and initial fraction of remote versus recent LTBI. We performed 1000 runs, so the 95% interval was obtained by taking the 25^th^ and the 975^th^ value in the list of outputs ordered from smallest to largest.

## Results

### Maximum likelihood curves

The best fit curves of the SEIS model to the number of PTB and EPTB cases per year are represented in Fig 2. Yearly immigrant numbers from the countries concerned are also represented (Fig 2, blue line). PTB cases have, on the whole, declined over the period 1995-20013, for all three ethnicities (Fig 2A,C,E). The strongest decline was seen among Turkish (Fig 2C) and was lowest among Indonesians (Fig 2E). Overall, most TB cases were found among Moroccans, then Turkish, and finally Indonesians. EPTB cases also tended to decrease, although less strongly than PTB, among Moroccans and Turkish (Fig 2B, D). In Indonesians the pattern was different, EPTB yearly cases being rather stable between 1995-2003, but with a peak in 2004-2005 (Fig 2F). In Turkish, the decline in PTB was very strong in comparison with the decrease in EPTB; the best-fit curve to the very low number of yearly PTB cases after 2009 matches poorly (Fig 2 C), given that EPTB cases showed a slight increase in that period (Fig 2D).

**Fig 2.**
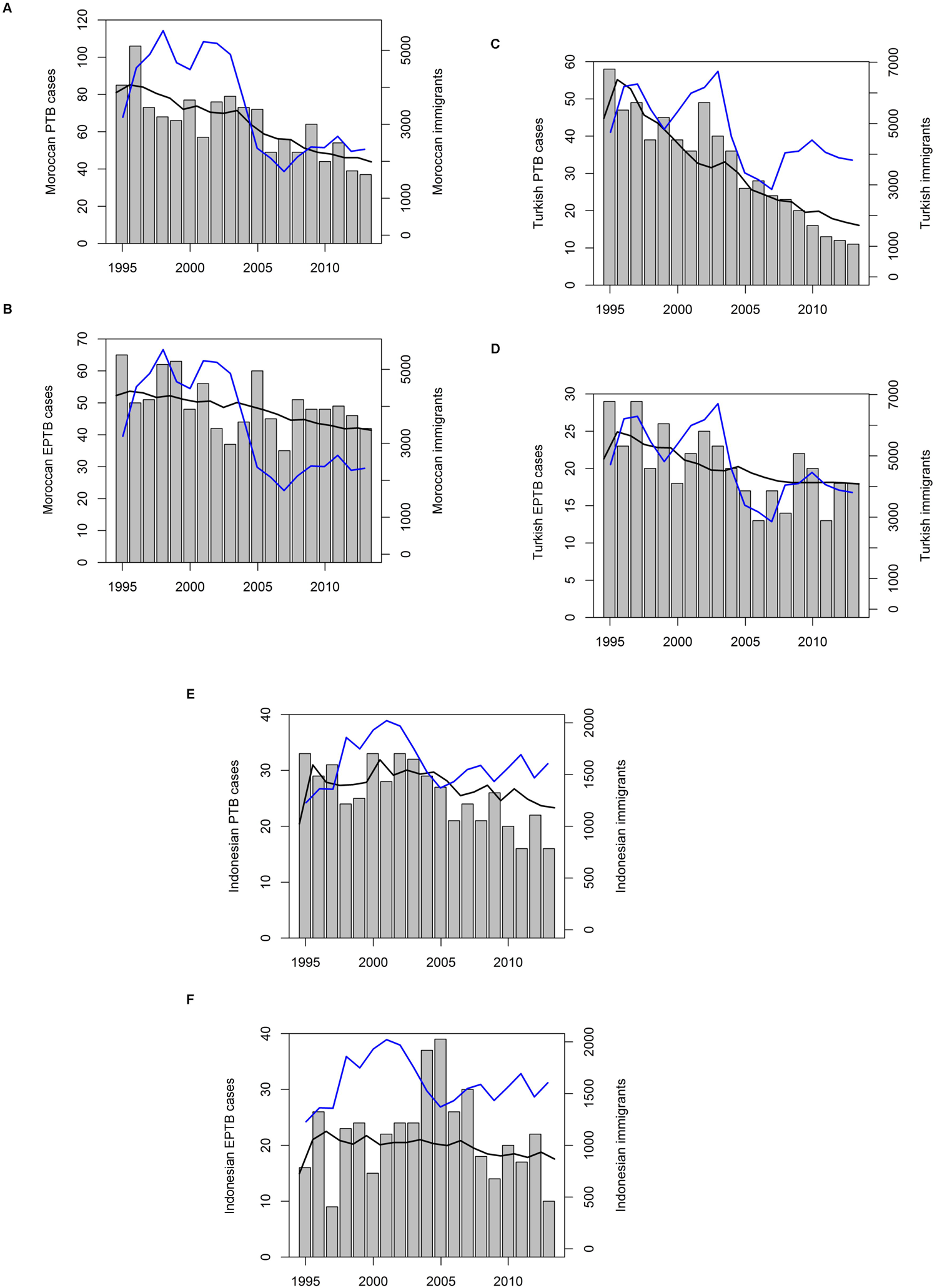
Best fit model to diagnosed (E)PTB cases. Number of diagnosed (A) PTB and (B) EPTB cases among Moroccan-born (grey bars; left axis). The black line represents the best-fit line to the diagnosed cases (left axis), the blue line represents yearly immigration figures from Morocco (right axis). Turkish-born (C) PTB, (D) EPTB cases, and Indonesian-born (E) PTB, (F) EPTB cases.

Immigration rates seem to have trends that did not match the number of TB cases observed. The exception is the number of yearly Turkish EPTB cases, which seem to follow roughly the line of immigrants (Fig 2D). The general pattern in immigration from Morocco and Turkey (Fig 2A and C) is first an increase from 1995 to 1997/98, a dip in 1999, then again a peak around 2002. A decline then follows, immigration being lowest in 2007. Then it picks up again, not quite reaching 1995 levels in 2013. Indonesian immigration figures were the lowest, with a different pattern over time (Fig 2E): there was a big increase from 1995 to 2001, followed by a decrease till 2001 to about 1995 levels, and then again a slow increase till 2013.

### Fitted parameters: transmission rates, fraction of recent versus remote infection and change in progression rate to EPTB vs PTB

Transmission rate was highest among Moroccans, with a rate *Ψ_M_* of 33/year (Table 2). Considering a PTB prevalence *P_PTB_M_* of about 25/100000 in the Moroccan community in the Netherlands (derived from best fit model predictions for PTB cases), this corresponds to 0.76% of Moroccan susceptibles becoming LTBI per year (*Ψ_M_*(1-*v_M_*))*p_PTB_M_*, so not taking into account travel to country of origin, see formula for *λ* in Methods). Transmission rate was about half the Moroccan transmission rate in the Turkish and Indonesian community (Table 2, 18/year and 14/year, respectively). Given a PTB prevalence of 10/100,000 in Turkish and 12/100,000 in Indonesians, this means that between 1995-2013, on average 0.16% of Indonesian and 0.17% Turkish became latently infected each year. The fraction of recent and remote LTBI among first-generation Moroccans was 2.2% and 23.2% respectively, as estimated by computing the area-under-the-curve for recent and remote LTBI and dividing by the sum of all Moroccan subpopulations (*S*+*I_1_*+*I_2_*+*E_1_*+*E_2_*). For first-generation Turkish, the fraction of recent and remote LTBI was 0.7% and 15.2% respectively and for first-generation Indonesians 0.6% and 24.6% respectively. The estimated fraction of recent LTBI (as opposed to remote LTBI) among all LTBI in 1995 was 15% in Moroccans, 11% in Turkish and 3% in Indonesians (Table 2). The proportion of LTBI developing PTB vs EPTB declined in Turkish and Moroccans, reflecting an increase in EPTB diagnosis over the study period compared with PTB diagnosis. In Indonesians, the slope of the change in fraction developing PTB vs EPTB was estimated at 5×10^−6^. We therefore proceeded with fitting a model not accounting for a change (results no shown).

**Table 2.**
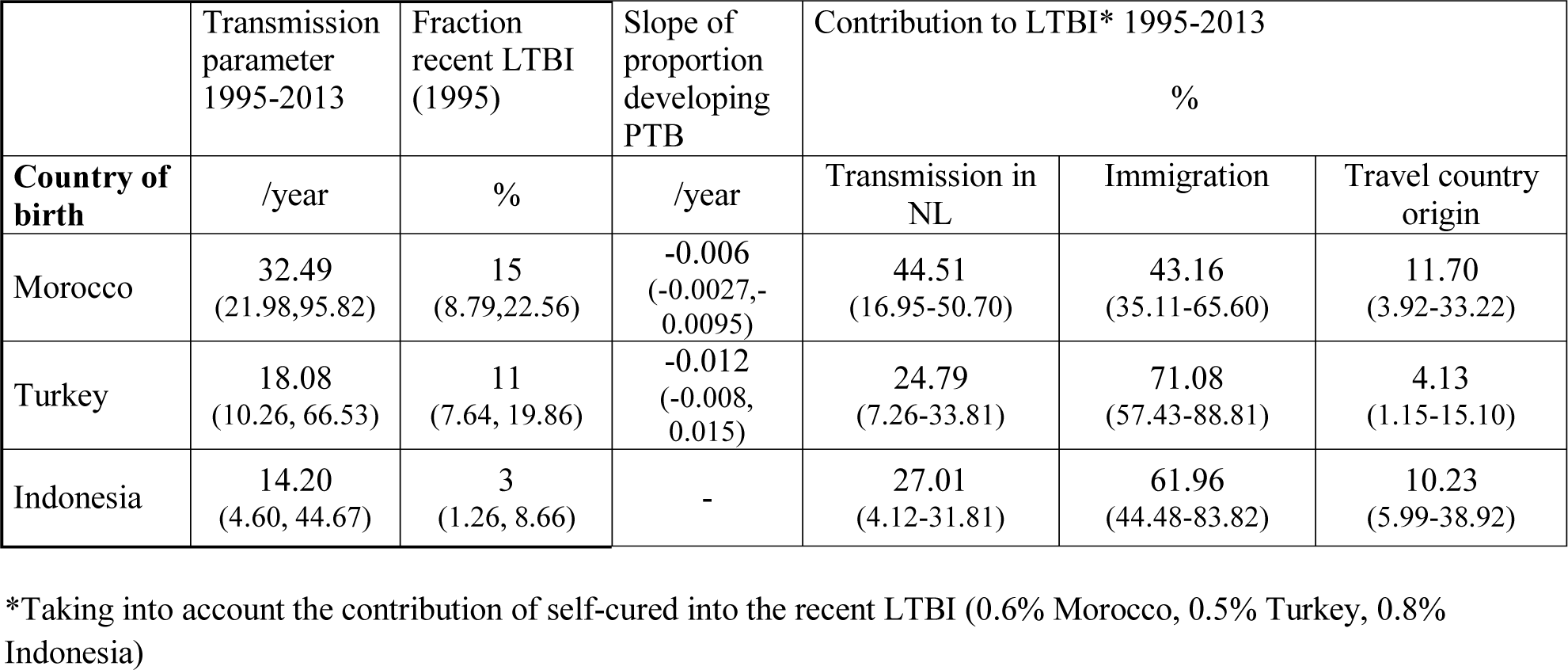
Estimates for transmission rate, fraction recent LTBI in 1995 and contribution of different routes to LTBI in the Netherlands for three ethnic groups –Morocco-, Turkey- and Indonesia-born communities. In brackets the 95% confidence regions of the multivariate sensitivity analysis.

|  | Transmission parameter 1995-2013 | Fraction recent LTBI (1995) | Slope of proportion developing PTB | Contribution to LTBI* 1995-2013 % |  |  |
| --- | --- | --- | --- | --- | --- | --- |
| **Country of birth** | /year | % | /year | Transmission in NL | Immigration | Travel country origin |
| Morocco | 32.49<br>(21.98,95.82) | 15<br>(8.79,22.56) | -0.006<br>(-0.0027,-0.0095) | 44.51<br>(16.95-50.70) | 43.16<br>(35.11-65.60) | 11.70<br>(3.92-33.22) |
| Turkey | 18.08<br>(10.26, 66.53) | 11<br>(7.64, 19.86) | -0.012<br>(-0.008, 0.015) | 24.79<br>(7.26-33.81) | 71.08<br>(57.43-88.81) | 4.13<br>(1.15-15.10) |
| Indonesia | 14.20<br>(4.60, 44.67) | 3<br>(1.26, 8.66) | - | 27.01<br>(4.12-31.81) | 61.96<br>(44.48-83.82) | 10.23<br>(5.99-38.92) |

\*Taking into account the contribution of self-cured into the recent LTBI (0.6% Morocco, 0.5% Turkey, 0.8% Indonesia)

The decrease in proportion LTBI developing PTB was strongest in the Turkish population: PTB numbers decreased much more strongly than EPTB cases over the period 1995-2013 (compare Fig 2D & 2E). This effect was seen in the Moroccan cases, but not as strongly (compare Fig 2A & Fig 2A & 2B); the yearly decrease in proportion LTBI developing PTB was twice as low in Moroccans (Table 2: −0.006/yr) compared to Turkish (-0.012/yr). In Indonesians, no consistent pattern could be fitted; this reflects the fact that EPTB cases showed a peak around 2005 (Fig 2F), while PTB cases remained constant initially, and decreased after 2005 (Fig 2E). Possible factors contributing to this pattern, such as the gradual ageing of TB cases in these groups over the study period, are discussed in the appendix (Section A4, Fig A1 and Fig A2).

Over most of the 1995-2013 period, transmission in the Netherlands and remote LTBI from immigration contributed roughly equally to LTBI in Moroccans (around 1000/year in 1995-2000, declining to about 500/700 in 2012-2013, Fig 3A). Travel contributed much less, starting at 300/year in 1995, and declining to about 200/year at the end of the period (Fig 3A). The smallest contribution was imported recent LTBI (Fig 3A). For the Turkish, the picture was slightly different, as imported remote LTBI was the most important contributor (around 800/year in 1995-2000, declining to around 550/year in 2013; Fig 3B). Second came transmission in the Netherlands: starting at 400 LTBI/year, it ended in 2013 at around 200 LTBI/year (Fig 3B). For Turkish, travel to the country of origin contributed least, lying just below the import of recent LTBI (about 50-100 LTBI /year; Fig 3B). The pattern in Indonesian LTBI contribution was very similar to that of the Turkish, but with a total absolute contribution that was half the contribution of the Turkish (about 600 total Indonesian LTBI /year, Fig 3C, compared to 1200 Turkish LTBI/year in the first period post-1995, Fig 3B).

**Fig 3.**
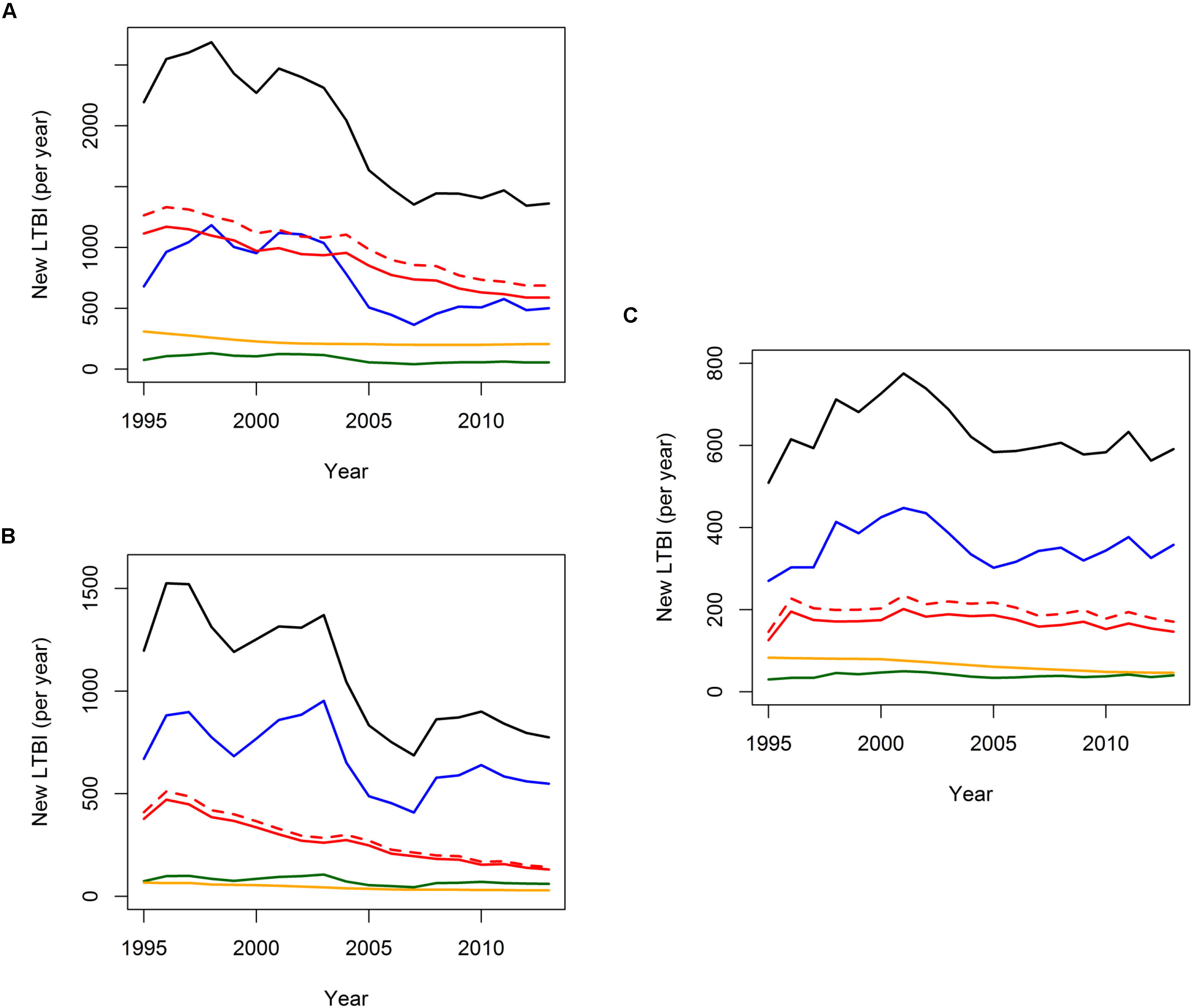
Model-derived yearly LTBI. Yearly new LTBI in the Netherlands among Moroccans (A), Turkish (B) and Indonesians (C): total LTBI (black), import of LTBI (remote LTBI: blue; recent LTBI: green), transmission in the Netherlands (red; including reinfection of remote LTBI: dashed red), travel to country of origin (yellow).

In the Moroccan community, we estimated that transmission within the Netherlands and immigration contributed almost equally to LTBI: 45% of new LTBI resulted from transmission in the Netherlands (excluding those resulting from reinfection in remote LTBI), while 43% resulted from immigration. Travel contributed more than in the other two communities, 12% (Table 2). In Turkish and Indonesians, similar percentages of LTBI came in through immigration (71% and 62% respectively). Transmission in the Netherlands contributed 25% and 27% of LTBI in Turkish and Indonesians respectively (Table 2). Travel contribution was lowest, namely 4% and 10% in Turkish and Indonesians respectively (Table 2).

The multivariate sensitivity analysis revealed broad 95% intervals for the transmission parameter (Table 2). The percentage of LTBI contributed by transmission within the Netherlands and by immigration had overlapping 95% intervals for Moroccans, but a significantly greater proportion of Moroccan LTBI arose from transmission in the Netherlands or came through immigration, than through travel to country origin. For Turkish and Indonesians, immigration contributed significantly more to LTBI than travel and transmission within the Netherlands (Table 2). The univariate sensitivity analysis, represented through a tornado plot (Appendix, FigA3), indicated that for all three countries, recovery of PTB, progression to 14 disease, reactivation rate and the extent to which transmission might be reduced in the Netherlands compared to the country of origin, are the parameters influencing TB transmission most. For the contribution to LTBI, the percentage attributable to transmission in the Netherlands for all three ethnicities was most affected by recovery through treatment, progression to disease, travel rate to country of origin, immune stabilization –*i.e.* transition from recent to remote LTBI- and reactivation (between about 5-13% deviation from estimate). Progression to disease, immune stabilization and reactivation rate influenced the percentage contribution to LTBI from immigration for all ethnicities (between 10-15% deviation from estimate). Recovery from PTB through treatment, travel to country of origin, and fraction reduction in transmission compared to country of origin were the most important parameters affecting the percentage contribution of travel to LTBI (travel to Turkey was least variable, with deviations below 5%).

## Discussion

The aim of our study was to estimate, based on a dynamic transmission model of TB in a low-incidence country, the size of the LTBI pool for three immigrant communities contributing most TB cases. Also, we derived the share of transmission within the Netherlands, versus immigration from and travel to country of origin, in the LTBI pool for these ethnic groups. We estimated that about 25% of Moroccans and Indonesians in the Netherlands are latently infected, of which 2% and 0.6% recently –in our definition less than 5 years ago-respectively. 16% of Turkish have LTBI, of which 0.7% recent. This may be set against results from Houben and Dodd (1), who provide estimates for the latent TB pool by world region: South East Asia (which includes Indonesia) has highest LTBI burden, with 30.8% prevalence; next is the Eastern Mediterranean region (Morocco) with 16% prevalence, and the European region (Turkey) with 13.7% prevalence.

Overall, immigration was the most important source of LTBI in the country: about 60-70% of LTBI among Turkish and Indonesians was due to immigration, significantly more than transmission within the Netherlands. For Moroccans, transmission within the Netherlands was equally important as immigration, roughly 45%.Travel to country of origin contributed little LTBI (at most 12%, for Moroccans).

Novel was the fit of our mathematical model to both PTB and EPTB incidences; as EPTB increased more than PTB diagnoses in Turkish and Moroccans, we had to account for an increasing ratio of EPTB *vs.* PTB diagnoses over time. This may reflect a shift to diagnoses of older TB cases in these communities, as EPTB is more often diagnosed in elderly. Yet, while the mean age at diagnosis did increase over the study period for Turkish and Moroccans, the association between age and site of disease was only apparent for Turkish. Our model could not fit the peak in Indonesian EPTB diagnoses in 2004-2005, and alternative hypotheses would have to be proposed to explain this observation: possibly, immigrant EPTB cases were diagnosed with a delay –absent for PTB cases- following the Indonesian immigration peak in 2002-2003 (23). The steady relative increase in EPTB diagnoses has been observed elsewhere -in the UK (24), in Spain (25) and in Serbia (26), and a more detailed epidemiological study would be interesting to shed light on the factors driving this relative increase.

As in Dowdy *et al.* (6), the univariate analysis shows that progression rate and recovery of disease are the parameters that influence the results most. Uncertainty in the travel rate, especially for Indonesia, can make a big difference in outcome, up to a 13% shift in percentage contribution to travel or transmission in the country. Research to pin down travel frequency would help to remove uncertainty in outcome.

We assumed random mixing within ethnic groups, and no mixing outside the ethnic groups. Borgdorff *et al.* (10), using TB fingerprinting data from all cases in the Netherlands between 1993-1995, showed that for Turkish, the assumption that most transmission occurs within the ethnic community is not unrealistic: of all Turkish cases, 20% result from infection by a Turkish source and 2% from another nationality, the rest being imported or source in a cluster. For Moroccans this is less pronounced, these percentages being 17 and 6% respectively (10). If we allowed for infectious contacts outside the ethnic group, there would be more potential for transmission -for example Moroccans could infect other ethnic groups, and Moroccans could get infected by other ethnicities. Therefore, we have provided an overestimate for the transmission within the ethnic group.

We assumed two categories of LTBI, a recently and a remotely infected group. We considered that recent infections, less than 5 years ago, progress faster to disease, while remote infections stabilize in latency with a very low probability of reactivating. However, the probability of progressing is much higher in the first 2 years than in the 3 years thereafter (27), and other modelling studies have used a 2-year cut-off (1). We don’t expect an impact of the progression rates of recent LTBI on the share of import versus transmission in the country. Only if progression rates differed between immigrants and residents would we expect an impact on the percentage contribution to LTBI.

The number of imported active TB cases was taken to be the cases diagnosed within 6 months of arrival in the country. This time was recorded for about 85-90% of cases. We essentially assumed that time since arrival in the Netherlands was always known when this was recent, and neglected those whose date of entry was not recorded. If recently immigrated cases were equally likely not to have a date of entry in the country recorded, we would have missed 14% and 11% of cases in Moroccans and Indonesians, respectively, *i.e.* by a relatively small percentage.

We assumed that 10% of the LTBI immigrating are recent infections. Literature on this percentage is scant, but Houben *et al.* have estimated that recent infection prevalence in the different world regions ranges from 1.2% in South East Asia, down to 0.3% in the European region, for an LTBI prevalence of 30.8% down to 13.7% respectively (1). Using this fraction as a lower estimate –as they concern infections that occurred less than 2 years ago, compared to 5 years in our model– we obtained a transmission parameter of 34/yr for Moroccans, 20.9/yr for Turkish, and 15.4/yr for Indonesians. Percentage contributions for transmission in the Netherlands change only slightly as a result: 45.5% for Moroccans, 27.4% for Turkish and 28.6% for Indonesians.

We have set out a framework to fit a dynamic transmission model to PTB and EPTB cases over a 20-year period, in order to quantify the contribution of transmission versus immigration and travel to LTBI. We have deliberately simplified the model to allow the fit to incidence data. We could envisage a number of features that may render the model more realistic.

First and most importantly, we would like to introduce the possibility of mixing between ethnic groups. Using the information from TB typing -MIRU-VNTR for culture-positive cases, and whole genome sequencing currently being developed- we would derive the transmission rates between groups. We have developed software as an add-on to BEAST (phylogenetic software package) that may help us quantify this transmission (Dhawan *et al.* in prep).

Second, age-structure could be added to the model, as it is known that rates of disease progression vary according to age.

Third, the information on fingerprinting could be used to better calibrate the model in terms of recent transmission versus reactivation cases. Unique fingerprints in immigrants who entered the country more than 2 years ago could be considered as a reactivation. An additional possibility would be to take into account a peak in progression rate at the time of immigration. Increased progression rates -due to high stress levels at immigration and mediated by increased cortisol levels, or reduced levels of vitamin D as a result of changing UV-B exposure- have been suggested (28, 29).

Finally, an important group of immigrants with high TB incidence consists of asylum seekers. As their stay in the country is not guaranteed, deriving estimates for this group requires a different modeling approach: it should be modelled as a separate entity, with probably little contact with other communities in the Netherlands as asylum seekers reside in centers, and with greater probability of leaving the country after a short period than immigrants with a residence permit.

In summary, we have formulated a dynamic model for TB transmission in a low-incidence country, accounting for import of cases from high-incidence areas. This model was fitted to (E)PTB incidence data for three ethnic groups. We showed that for these ethnic groups, travel to country of origin contributed least to new LTBI, and immigration was the most important contributor. Only for Moroccans was the share of transmission in the Netherlands equally important as immigration. These findings have a implications for policy scenarios: targeting returning travelers might be less effective at preventing LTBI than screening for LTBI at entry in the country. Also, contact investigation might well have different yields between ethnic groups (5), as transmission rates among Moroccans would appear to be higher than for other ethnicities.

